# Integrating genomic and tagging data reveals spatio-temporal population structure in Northeast Atlantic European sea bass

**DOI:** 10.64898/2026.06.22.731647

**Authors:** Pierre-Alexandre Gagnaire, Mathieu Woillez, Hélène de Pontual

## Abstract

Understanding spatial and temporal connectivity among individuals with different migration strategies is essential for migratory ecology and effective conservation, yet it often requires integrating multiple data sources. In Northeast Atlantic European sea bass (*Dicentrarchus labrax*), electronic tagging has revealed partial migration, with both resident and long-distance migrants showing fidelity to summer feeding and winter spawning areas. However, the role of regional spawning-site philopatry in shaping migration patterns and stock connectivity remains unclear. Here, we combine reconstructed migration trajectories with genome-wide analyses of gene flow and recent relatedness in 708 individuals sampled from 10 French Atlantic locations. We identify a seasonally shifting genetic discontinuity between the Bay of Biscay (BOB) and Northern (NS) stocks, located off western Brittany during winter spawning and displaced northeastward into the central English Channel during summer feeding. Despite seasonal mixing in the English Channel, an association between individual genetic composition and spawning-site selection supports regional spawning-site philopatry. Analyses of long genomic segments shared identical-by-descent reveal substantially greater connectivity within stocks than between stocks, indicating that philopatry constrains effective gene flow despite seasonal mixing. Reanalysis of independent genomic data further shows that sea bass from the northern Atlantic range predominantly belong to the Northern stock. Together, these results show how seasonal movements reshape spatial genetic structure while maintaining demographic subdivision, with direct implications for fisheries management.

## Introduction

Population structure arises from interactions among geographic, demographic, and behavioral factors that lead to non-random spatial assortment of individuals during reproduction (Slatkin, 1987). Understanding population structure therefore provides key insights into the demographic and biological processes shaping mating patterns and population connectivity (Lowe & Allendorf, 2010). This knowledge is fundamental for identifying and delineating the demographic units that require conservation and management monitoring (Lande, 1988).

Mobile species with complex migratory life cycles pose particular challenges for connectivity studies. In these species, recurrent migratory movements continually reshape the spatial distribution of individuals, resulting in population structure that varies over time. This situation is typical of species that alternate between breeding and foraging habitats, sometimes including overwintering areas, connected by long-distance migrations. Studies in cetaceans (Hoelzel, 1998), sea turtles (Bowen et al., 2005), migratory birds (Ruegg et al., 2014) and insects such as monarch butterflies (Freedman et al., 2021) have addressed these challenges, revealing key aspects of complex life cycles that are often inaccessible through direct observation. Many marine fish species that undertake seasonal or terminal spawning migrations between feeding and spawning habitats face similar difficulties.

Despite being hidden beneath the ocean surface, migration patterns of marine fishes have been increasingly documented using electronic tagging techniques (Block et al., 2026). Unlike terrestrial species, marine mammals, or sea turtles that can be tracked via satellite positioning, fish movements cannot be directly geolocated because seawater does not transmit the electromagnetic signals required for GPS. Instead, positions are reconstructed from data recorded by acoustic or archival tags (Aarestrup et al., 2009; Block et al., 2005). Advances in Data Storage Tags (DSTs) now allow long-term, high-frequency recording of environmental variables such as temperature, pressure, and light. Geolocation models can then infer individual migration trajectories from these environmental time series (Braun et al., 2018; Woillez et al., 2016). Such approaches can detect site fidelity and temporary mixed assemblages of demographic units in shared feeding areas, overlapping migratory routes or common wintering areas (Hunter et al., 2003; Neuenfeldt et al., 2013). However, while reconstructed trajectories describe individual movement and spatial overlap, they lack information on how migration patterns translate into genetic exchange among groups of individuals.

Genetic approaches offer a complementary perspective by informing on migration rates and gene flow (Hedgecock et al., 2007; Lowe & Allendorf, 2010). Spatial genetic structure can be used to delineate stocks and conservation units, estimate levels of connectivity, and assess the roles of adult philopatry and larval dispersal (Cayuela et al., 2018; Funk et al., 2012; Hohenlohe et al., 2021). Nevertheless, traditional population genetic analyses generally reflect long-term evolutionary processes and may fail to capture seasonal movements and their impact on spatial genetic patterns. Moreover, static genetic sampling designs are ill-suited to represent the dynamic population structure of migratory species unless sampling is explicitly temporally resolved. Integrating spatial genetics with spatio-temporal tagging data therefore offers strong potential to resolve connectivity dynamics while reducing mismatches in temporal scale (Delgado et al., 2025; Hawkins et al., 2016; Kool et al., 2013; Mikles et al., 2026). Here, we apply this strategy to investigate spatio-temporal connectivity in European sea bass populations of the Northeast Atlantic.

The European sea bass (*Dicentrarchus labrax*) is a highly valued species in the Northeast Atlantic, exploited by both recreational and commercial fisheries. Its distribution ranges from the Scottish and Norwegian coasts in the north to Morocco, the Mediterranean Sea, and the Black Sea in the south (Vandeputte et al., 2019). Sea bass rely on distinct habitats throughout their life cycle (López et al., 2015; Pickett & Pawson, 1994). Reproduction occurs offshore, after which eggs and larvae are advected toward estuarine or sheltered coastal nursery areas that support juvenile growth and survival. Upon maturity, adults undertake seasonal migrations between coastal feeding areas in summer and offshore spawning grounds in winter.

Historically, European sea bass fisheries were managed at the national level until a sharp decline in stock biomass during the 2010s prompted the introduction of emergency conservation measures by the European Union (EU, [2015/2596(RSP)](https://eur-lex.europa.eu/legal-content/EN/TXT/?uri=CELEX:52015IP0078)). These measures rely on management units, or stocks, defined by the International Council for the Exploration of the Sea (ICES). Catch reductions were recommended for the “Northern stock” (NS), encompassing the Irish Sea, Celtic Sea, English Channel, and southern North Sea, and separated from the “Bay of Biscay stock” (BOB) by the 48th parallel north. Two additional stocks have been delineated: one extending from southern Ireland to western Scotland, and another off the coasts of Portugal and northern Spain (**Fig. 1A**).

**Figure 1.**
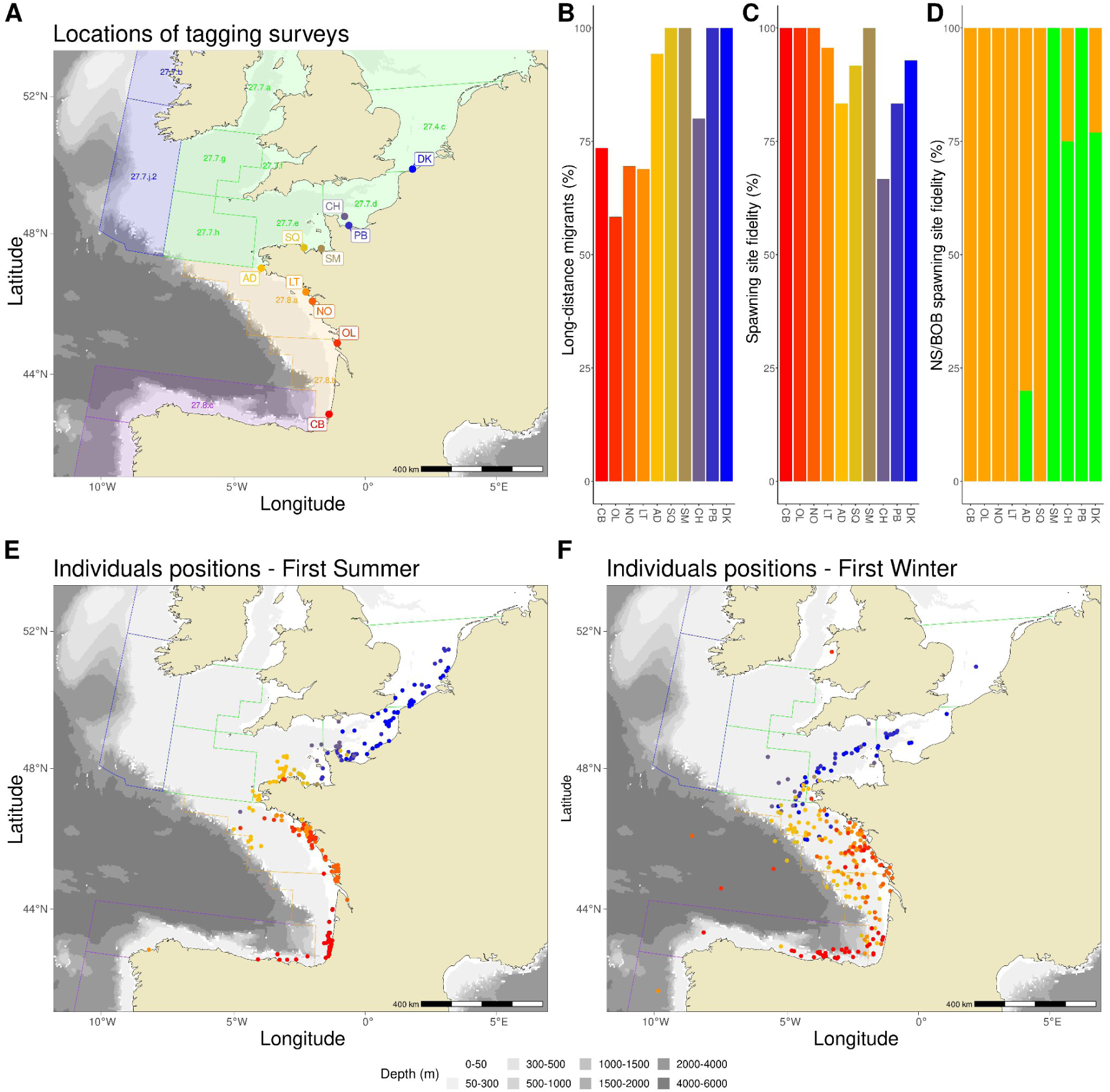
Tagging locations and seasonal migration patterns of adult European sea bass. (**A**) Locations of the 10 tagging sites along the French Atlantic coast. Shaded regions indicate the four ICES management units for European sea bass in the Northeast Atlantic: west of Ireland and Scotland (blue), Northern stock (NS; green), Bay of Biscay stock (BOB; orange), and Iberian stock (purple). For each tagging site, (**B**) the proportion of migratory individuals among all tagged fish classified as resident or migratory (N=248), (**C**) the proportion of migrants exhibiting spawning-site fidelity (N=112), and (**D**) the relative proportions of migrants showing fidelity to NS (green) or BOB (orange) spawning grounds (N=103) are shown. (**E**) Average individual positions during the first summer feeding season following tagging (N=320). (**F**) Average individual positions during the first winter spawning season following tagging (N=248).

In successive assessments of the NS and BOB stocks (ICES, 2025a, 2025b), ICES experts have emphasized the need for a better understanding of spatial and temporal population structure. Such knowledge is essential to avoid mismanagement arising from stock boundaries that do not reflect self-sustaining discrete populations (Cadrin, 2020; Cadrin et al., 2023; Kerr et al., 2017). Mark-recapture and archival tagging studies have shown that European sea bass exhibit partial migration, with some individuals remaining resident while others migrate over long distances between summer feeding areas and offshore winter spawning grounds (de Pontual et al., 2019; de Pontual et al., 2023; Goossens et al., 2024; Wright et al., 2024). Although strong interannual spawning-site fidelity has been documented (de Pontual et al., 2019; de Pontual et al., 2023; Le Luherne et al., 2022), most population genetic studies have reported little to no spatial genetic structure across the Northeast Atlantic (Coscia & Mariani, 2011; Fritsch et al., 2007; Souche et al., 2015).

A population genomic study with broad spatial coverage leveraged the Mediterranean ancestry gradient along the Atlantic coast to identify two subtle genetic discontinuities, likely separating demographically independent regional stocks (Robinet et al., 2020). One of these discontinuities was detected in northern Brittany, near the Gulf of Saint-Malo, and may correspond to the boundary between the NS and BOB management units. By contrast, a more recent study based on the sea bass 57k SNP array (Griot et al., 2021), found no evidence for genetically differentiated populations within the northern Atlantic range, concluding that the NS and BOB stocks do not constitute distinct populations (Taylor et al., 2025). Differences in spatial and temporal sampling resolution between these studies may partly explain these contrasting conclusions. Together, these results indicate that the extent to which regional philopatry limits gene flow between the NS and BOB stocks, and how seasonal migrations influence the spatio-temporal dynamics of this boundary, remains unresolved.

In this study, we use the sea bass 57k SNP array to conduct a high-density genomic analysis of connectivity in European sea bass sampled as part of a large-scale electronic tagging program that tracked individual migration trajectories over two consecutive years (de Pontual et al., 2023). We examine spatial genetic structure at two key stages of the annual cycle: summer, when adults occupy feeding habitats, and winter, when reproduction occurs on spawning grounds. By explicitly integrating genomic and movement data across seasons, we reveal the spatio-temporal dynamics of a shifting connectivity boundary between the NS and BOB stocks in the western English Channel, with direct implications for the management of European sea bass.

## Materials and Methods

### Individual electronic tagging and recapture

Samples used in this study were collected as part of a large-scale electronic tagging programme (de Pontual et al., 2023). Briefly, ten electronic tagging surveys were conducted along the French Atlantic and English Channel coasts during the summers 2014, 2015, and 2016 (**Suppl. Table S1**, **Fig. 1A**). Adult sea bass were equipped with data storage tags (DSTs) following a tagging protocol described in (de Pontual et al., 2019). During tagging, caudal fin clips were collected from each individual and preserved in 95% ethanol for subsequent genetic analysis. Given the two-year battery life of the DSTs, tags were programmed to record temperature and pressure (i.e., depth) every 90s for the first 680 days, and every 3min for the remaining 50 days. In total, 1,220 adult sea bass were tagged, of which 482 individuals (39.5%) were recovered by November 2022 through either fish recapture or tag recovery. Recoveries were relatively evenly distributed between the Bay of Biscay (BOB) and Northern Stock (NS) areas.

### Reconstruction of individual trajectories

Temperature and depth time series recorded by DSTs were used to reconstruct individual fish trajectories using a geolocation Hidden Markov Model (HMM) (Thygesen et al., 2009), adapted from previous studies on sea bass (de Pontual et al., 2019; Woillez et al., 2016). In this model, daily fish position is treated as the unobserved state of a dynamic system governed by a behavioral switching motion model (similar to Pedersen et al., 2008) and an observational model linking environmental reference fields to the recorded depth and temperature series (de Pontual et al., 2023). The movement model is based on a random walk with two behavioral states - high and low activity - identified from individual vertical movement patterns (Heerah et al., 2017).

Model outputs consist of daily posterior probability distributions representing uncertainty in estimated positions. Individual trajectories were then derived as the most probable sequence of daily locations. Trajectories were first compiled per tagging site to characterize post-tagging migration strategies (de Pontual et al., 2023). Individuals were classified as residents if they remained within 50 km of their tagging location between their first summer and winter locations. Fish moving beyond this range, particularly during winter, were categorized as migrants.

Site fidelity was defined as the tendency of migrants to return to the same functional habitat. Feeding-ground fidelity was identified when individuals returned to forage within 50 km of their tagging location during two consecutive summers. Spawning-ground fidelity was defined as return to the same ICES stock area (NS or BOB) for reproduction during two consecutive winters. Differences in migration strategies between NS and BOB stocks were assessed using binomial generalized linear models (GLMs) fitted in R, with tagging sites treated as replicates.

To evaluate how adult migrations and site fidelity influence genetic structure, individual trajectories were summarized to represent seasonal occupancy of key functional areas. Since tagging surveys were conducted between June and early September (**Suppl. Table S1**), the first summer distribution of individuals in coastal and inshore feeding habitats was represented by the average position during the last two weeks of September in the year following tagging. Adult sea bass exhibit increased vertical activity during the spawning season (Heerah et al., 2017). Therefore, spawning distribution was represented by the average positions corresponding to the highest two-week mean depths during the spawning months (January-March) of the first winter after tagging.

### SNP Genotyping using the Axiom DlabChip array

Whole genomic DNA was extracted following proteinase K digestion using a standard protocol, and DNA quantity and quality were assessed with a Nanodrop spectrophotometer. Genotyping was performed by the Gentyane INRAe platform (Clermont-Ferrand, France) using the ThermoFisher Axiom™ Sea Bass 57k SNP *DlabChip* (Griot et al., 2021). This high-density SNP array was designed to capture genome-wide variation across the European sea bass genome, with markers evenly distributed with respect to recombination distance. Marker selection also aimed to enrich for intermediate-frequency variants across the species’ range, including both Atlantic and Mediterranean lineages, in order to minimize ascertainment bias and maximize power for ancestry inference.

In total, 768 sea bass were genotyped using two 384-well plates of the Axiom *DlabChip* array. This dataset comprised 708 spatially balanced individuals from the 10 tagging sites along the French Atlantic and English Channel coast, ensuring balanced representation of the BOB and NS stocks. Among these, 482 individuals were recaptured, and 364 had trajectories successfully reconstructed from electronic tagging data. An additional 60 individuals from the Algarve region (Faro, FA) in southern Portugal were genotyped to serve as a reference population characterized by elevated Mediterranean ancestry (Robinet et al., 2020). This reference was used to facilitate the detection of Mediterranean introgression signals within northeastern Atlantic sea bass populations.

Genotype calling and quality control were performed using *Axiom Analysis Suite v3.1.51.0* (Affymetrix). To enhance the detection of low-frequency Mediterranean alleles in Atlantic samples, the two Atlantic plates were analyzed jointly with a third plate containing Mediterranean individuals and experimental Atlantic-Mediterranean hybrids from an independent study. Plate-level quality control (QC) thresholds were set as follows: DQC ≥ 0.82, QC call rate ≥ 90%, passing samples ≥ 95%, average call rate for passing samples ≥ 98.5%. SNP classified as OTV, MonoHighResolution, CallRateBelowThreshold and Other were excluded. Additional SNP-level filtering included a minor allele frequency (MAF) threshold of 1% and the removal of SNPs deviating from Hardy-Weinberg equilibrium (HWE) within northeast Atlantic samples (excluding Portuguese reference individuals), using a P-value threshold of 0.0001. This HWE filter was applied to remove loci affected by genotyping artefacts while retaining markers reflecting true population structure.

### Analysis of population genetic structure

Population genetic structure of Northeast Atlantic European sea bass was assessed using a final dataset of 47,211 SNPs polymorphic within Atlantic populations. This dataset included high-quality genotypes for 705 individuals sampled along the French Atlantic and English Channel coast, as well as 60 individuals from the Algarve region (Portugal). Genetic diversity was quantified for each of the 11 sampling locations using mean observed (*H*_o_) and expected (*H*_e_) heterozygosity.

Pairwise genetic differentiation among sampling locations was assessed using *F*_ST_ (Weir & Cockerham, 1984), as estimated by the *StAMPP v1.6.3* R package (Pembleton et al., 2013). Confidence intervals and P-values were obtained from 10,000 bootstrap replicates across loci. To identify population genetic clusters and assign membership of tagging sites, hierarchical clustering was performed on the pairwise *F*_ST_ matrix using the *hclust* function in R. Fine-scale population structure was further examined using Principal Component Analysis (PCA) conducted with the *adegenet v2.1.3* R package (Jombart & Ahmed, 2011).

To quantify spatial variation in Mediterranean introgressed ancestry, individual ancestry proportions were inferred using *Admixture* (Alexander et al., 2009), assuming two ancestral populations (*K*=2). The termination criterion for the optimization algorithm was set to 100 iterations, and 1000 bootstrap replicates were used to estimate the standard errors of individuals ancestry proportions. Mean Mediterranean ancestry and associated standard errors were then calculated for each tagging location and related to geographic distance from the Algarve reference population (Faro). Distances were computed as least-cost paths restricted to waters shallower than 200 m, using the R package *marmap* (Pante & Simon-Bouhet, 2013). The relationship between Mediterranean ancestry and distance was modeled using a transformed logistic-shaped hyperbolic tangent (*tanh*) function fitted with *nls* in R, a commonly applied approach for modeling geographic clines.

### Estimation of effective migration surfaces integrating reconstructed trajectories

When dispersal is spatially restricted, genetic similarity typically declines with geographic distance, resulting in isolation-by-distance (Wright, 1943). Local reductions in gene flow, arising from physical or behavioral barriers, can generate abrupt increases in genetic differentiation that depart from this general pattern. Such spatial variation in migration rates can be quantified using the method *eems* (Petkova et al., 2016), which relates genetic similarity to both local and global migration rates to estimate effective migration surfaces.

We used a fast implementation of *eems* called *feems* (Marcus et al., 2021), to infer spatial patterns of genetic connectivity in European sea bass. A discrete spatial grid was constructed by partitioning the Earth’s surface into equally sized triangular cells using *dggridR v2.0.8* (Barnes, 2017). A dense triangular lattice covering the Northeast Atlantic was generated by defining a polygon envelope around sample locations and excluding land areas. Grid parameters were set to Projection = ISEA, aperture = 4, res = 8, and precision = 7, providing a compromise between spatial resolution and noise reduction.

The *feems* analysis was conducted using sample locations derived from reconstructed individual trajectories and custom parameter values (lamb_l2 = 0.0, lamb = 1e-8, alpha = 5). To explicitly account for seasonal migrations between feeding and spawning habitats, effective migration surfaces were estimated separately for summer (feeding) and winter (spawning) periods.

For the reproductive season, individual average positions during the first winter after tagging were used to characterize the spatial genetic structure of breeding units within their spawning areas. The resulting winter effective migration surface thus reflects genetic connectivity among reproducing groups and provides insight into breeding stock subdivisions.

For the feeding season, individual average positions at the end of the first summer were used to represent distribution within coastal and inshore feeding habitats. The corresponding summer effective migration surface was used to assess how feeding migrations modify the winter spatial genetic structure. Specifically, this analysis aimed to determine whether feeding site fidelity simply shifts the spatial location of genetic structure or leads to a more complex reorganization through the seasonal mixing of individuals from different spawning stocks within shared feeding areas.

### Test of regional philopatry

Regional spawning-site philopatry was tested within the English Channel, where stock mixing occurs during the summer feeding season (de Pontual et al., 2023). Because tagging was conducted during summer, we combined individual genetic data with reconstructed winter positions to assess whether spawning migration behavior was associated with genetic background, while controlling for variation among tagging locations. Analyses were restricted to individuals tagged within the English Channel for which winter positions could be reconstructed. The SM location was excluded due to sample size of one.

To test for a relationship between genetic composition and spawning-site selection, we fitted a linear mixed-effect model using the *lme4* R package (Bates et al., 2003). The model included both random-intercepts and random-slopes for tagging location, allowing baseline spawning latitude and the strength of the genetic effect to vary among locations. Individual genetic composition was quantified using scores on the first PC axis of the PCA, and spawning-site location was represented by mean latitude during the first winter after tagging.

To assess whether spawning latitude might be influenced by loci with intermediate to large effects, we further conducted a Genome-Wide Association Study (GWAS) using the R package *statgenGWAS* (Van Rossum & Kruijer, 2020). Individual mean winter spawning latitude was treated as a continuous phenotypic trait.

### Analysis of recent stock connectivity through IBD sharing

We assessed recent genetic connectivity among BOB and NS sea bass stocks by identifying long genomic segments identical-by-descent (IBD) using *IBIS v.1.20.6* (Seidman et al., 2020), which infers IBD segments and pairwise kinship coefficients from unphased genotype data. Analyses were performed using the following parameters: -min_l = 7, -a = 0, -maxDist = 0.1, -mt = 100, -er = 0.004, -ibd2, -mt2 = 50, and -printCoef. We focused on shared segments longer than 7 cM, a threshold corresponding to the highest IBD detection accuracy reported for IBIS. IBD segment lengths were expressed in recombination units using a high-density genetic map developed for Atlantic European sea bass families genotyped with the same SNP array (Leitwein et al., 2024).

Pairwise kinship coefficients were averaged among individuals to construct a matrix of mean relatedness between tagging sites, which was then used for hierarchical clustering to identify clusters of recent genetic coancestry. To quantify recent connectivity between stocks, we estimated the proportion of related individual pairs within and between the BOB and NS stocks, retaining only pairs with non-zero kinship coefficients and a reported degree of relatedness equal to or closer than seventh degree.

Finally, to examine how recent relatedness varies with spawning distance, we analyzed pairs of individuals with reconstructed winter positions. Least-cost distances between winter spawning locations were calculated for each pair, binned into 19 quantiles containing equal number of observations, and used to compute mean kinship coefficients within each bin. Analyses were conducted separately for pairs of individuals from the same stock (BOB-BOB and NS-NS) and from different stocks (BOB-NS). For each pair type, the relationship between average kinship coefficient and spawning distance was tested using beta regression.

### Integration of external genomic data

To place our results in a broader geographic context, we integrated published genotype data from Atlantic European sea bass populations surrounding the UK that were generated using the same SNP array (Taylor et al., 2025). This dataset comprised 558 individuals sampled in feeding habitats across 20 locations and 286 individuals sampled in spawning habitats across 15 locations, genotyped at 41,022 SNPs. After merging with our dataset and applying consistent quality filtering, the combined dataset included 1,609 individuals genotyped at 39,122 SNPs.

We performed a PCA including all individuals from the Taylor’s dataset together with our 10 French tagging locations. Samples from southern Portugal were included to orient the first principal component (PC1) along the gradient of Mediterranean ancestry. Using individuals from our study with both genetic data and reconstructed spawning locations, we defined empirical PC1 distributions and their associated 95% confidence intervals for the BOB and NS spawning stocks using 1000 bootstrap replicates. The PC1 coordinate distribution of individuals from Taylor’s dataset was then compared to these reference distributions to infer mixture proportions with the approach detailed below.

### Inference of mixture proportions using reference baselines

Mixture composition was inferred at the sample level using an empirical Bayesian mixed-stock framework. Reference density distributions were estimated from baseline samples and subsequently used for Bayesian inference of mixture proportions without individual assignment. Reference samples from the two spawning stocks BOB and NS were defined based on winter latitude and used to estimate empirical kernel density distributions of individual PC1 coordinates. For each mixture sample, individuals were treated as independent draws from a two-stock mixture, with the likelihood of each PC1 value expressed as a weighted sum of the two reference densities. The proportion of the BOB stock was inferred by evaluating the posterior distribution of the mixture parameter over a fine grid between 0 and 1, assuming a uniform Beta(1,1) prior. The Maximum A Posteriori (MAP) estimate was retained as the point estimate, and posterior 95% credible intervals were used to quantify mixture sampling uncertainty. Spatial patterns of inferred mixture proportions were visualized by generating maps across ICES areas separately for feeding and spawning seasons, using individual PC1 coordinates derived from the analysis of merged datasets.

Sensitivity to baseline estimation was assessed by resampling reference individuals with replacement, re-estimating reference densities, and repeating mixture inference across 1000 bootstrap replicates. Uncertainty attributable to baseline estimation was summarized using percentile intervals derived from the bootstrap distribution of MAP estimates and is reported separately.

## Results

### Partial migration and spawning-site fidelity

A total of 364 individual sea bass trajectories were reconstructed (**Suppl. Table S1**). The difference between the number of recovered tags and reconstructed trajectories reflects computational limitations affecting a subset of tags. Building on previous analyses (de Pontual et al., 2023), we analyzed site-specific migration strategies and confirmed that European sea bass in the Northeast Atlantic exhibits a partially migratory meta-population structure (Reid et al., 2018), with both resident and migratory individuals, although migrants predominated. Fidelity to coastal and inshore feeding areas in summer and offshore spawning grounds in winter was observed across all sites.

Migratory individuals were significantly more frequent in the Northern Stock (NS) compared to the Bay of Biscay (BOB) area (**Fig. 1B**, binomial GLM P-value = 0.0002). In contrast, the proportion of migrants exhibiting spawning-site fidelity did not differ significantly between stocks (**Fig. 1C**, P-value = 0.185). Migrants with spawning fidelity used distinct breeding regions in the NS and BOB, consistent with regional spawning fidelity (**Fig. 1D**, P-value = 1.14e-6). At several tagging sites (AD, CH and DK), individuals tagged in summer migrated either southward to the Bay of Biscay or northward to the 48th parallel during winter, indicating localized stock mixing north of AD.

During the first summer after tagging, sea bass were broadly distributed from northern Spain to the southern North Sea, with localised feeding aggregations around estuarine and nearshore hotspots (**Fig. 1E**). In the subsequent winter, individuals from sites predominantly associated with an NS spawning strategy (SM to DK) migrated southwest from their summer locations (**Fig. 1F**, **Suppl. Fig. S1**), concentrating mainly in the western English Channel between the northern Cotentin Peninsula and the Iroise Sea. In contrast, individuals from sites primarily associated with a BOB spawning strategy (SQ to CB) moved southward within the Bay of Biscay (**Fig. 1F**, **Suppl. Fig. S1**), with some individuals from CB migrating westward along the northern Iberian coast. High winter densities at coastal sites near the Loire River and the Pertuis region largely reflected resident individuals undertaking short offshore spawning movements.

### SNP genotyping summary statistics

Quality control of the two genotyping plates resulted in the retention of 765 of the 768 genotyped individuals. The mean call rate for passing samples was 99.47% for the first plate and 98.77% for the second. Of the 56,730 markers on the array, 49,655 variants (87.5%) were classified as high-quality SNPs (PolyHighResolution), and 1,307 (2.3%) as rare variants lacking minor allele homozygotes (NoMinorHom). Five mitochondrial SNPs used to assign maternal lineage (Atlantic or Mediterranean) were also successfully genotyped (Hemizygous). After applying population-level filters (MAF > 0.01, HWE P-value > 0.0001) to the French Atlantic samples, excluding the Portuguese individuals to limit the influence of population structure, the final dataset comprised 47,211 high-quality SNPs across 765 individuals. The mean proportion of missing genotypes per individual was 0.43%, with an average heterozygosity of 0.312 and a mean *F*_IS_ of 0.0049.

### Barrier to gene flow between BOB and NS stocks

Pairwise genetic differentiation among the ten locations sampled along the French Atlantic coast was consistently low, with *F*_ST_ values ranging from 0 to 0.00052 (mean = 0.00019). In contrast, comparisons involving the Portuguese sample from Faro showed substantially higher differentiation, with *F*_ST_ values averaging 0.00546, nearly 30 times greater than those observed within the French Atlantic. Hierarchical clustering based on pairwise *F*_ST_ values identified three genetic clusters (**Fig. 2A**): Cluster 1 comprised southern Portugal (FA); Cluster 2 included BOB and western English Channel locations (from CB to SM); and Cluster 3 encompassed eastern English Channel NS locations (from CH to DK). Genetic homogeneity, reflected by non-significant *F*_ST_ values, was observed within Cluster 2 and within Cluster 3. In contrast, most pairwise comparisons between clusters were highly significant, with the exception of comparisons involving SM, likely due to its smaller sample size (N=15). The SQ sample was differentiated from both Clusters 2 and 3 and occupied an intermediate position between them.

**Figure 2:**
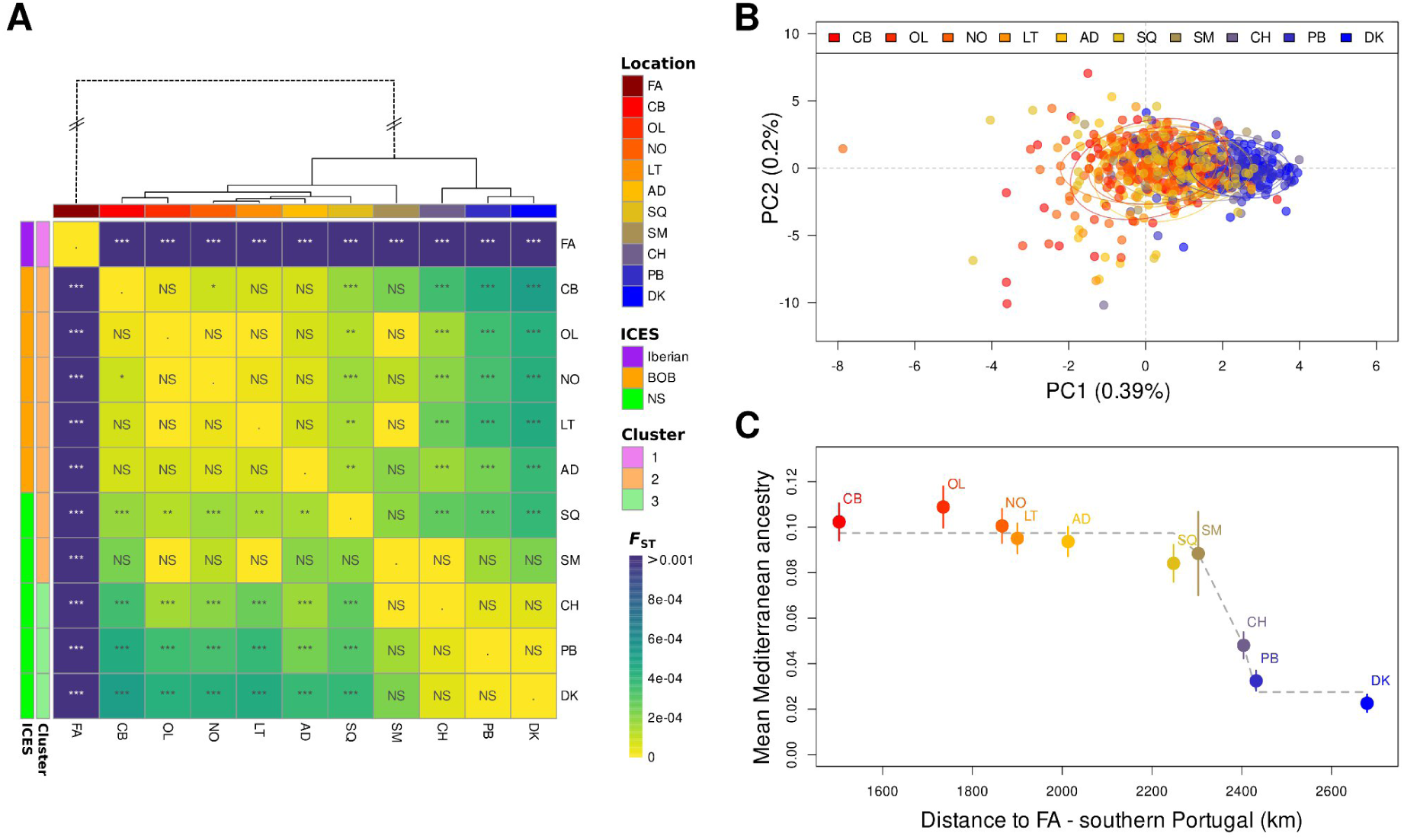
Population genetic structure of European sea bass along the French Atlantic coast. **(A)** Heatmap of pairwise *F*_ST_ values among tagging locations, organized by hierarchical clustering, revealing three genetically differentiated clusters. Significance levels are indicated as follows: NS (p > 0.05), ** (p < 0.01), and *** (p < 0.001). For visualization, the branch length separating the Faro (FA) sample from the French Atlantic locations was shortened (dotted branch). **(B)** Principal Component Analysis (PCA) highlighting the genetic discontinuity between Bay of Biscay (BOB) and Northern stock (NS) locations in the eastern English Channel. The first principal component (PC1) is oriented toward maximum differentiation from the Faro sample (see **Suppl. Fig. S2**) and captures the latitudinal gradient in Mediterranean ancestry, revealing a sharp transition west of the Cotentin Peninsula. **(C)** Spatial gradient in Mediterranean ancestry along the French Atlantic coast. Mean Mediterranean ancestry (± s.e.) is shown for each tagging location as a function of the least-cost distance to the Faro sample, calculated through waters shallower than 200 m. The dashed line represents the fitted geographic cline, highlighting the discontinuity between SM and CH.

The pronounced differentiation between southern Portugal and French Atlantic locations was captured by the first principal component (PC1) of the PCA, which reflected a latitudinal gradient in Mediterranean ancestry (**Suppl. Fig. S2**). Individuals from the Faro sample spanned a wide range of PC1 values, indicating substantial inter-individual variation in Mediterranean ancestry within this Atlantic-Mediterranean transition zone. This pattern is consistent with southern Portugal acting as a bridge for the introgression of Mediterranean alleles into the Northeast Atlantic. When focusing on the French Atlantic samples, PC1 further revealed finer-scale structure separating BOB and western English Channel locations from eastern English Channel NS locations, with a subtle but distinct spatial subdivision west of the Cotentin Peninsula (**Fig. 2B**). Notably, this discontinuity was evident only along PC1, highlighting its association with the Mediterranean ancestry gradient.

Individual ancestry estimates corroborated these results. Southern Portugal exhibited the highest Mediterranean ancestry, with individual proportions ranging from 0.10 to nearly 1 (mean = 0.61, s.d. = 0.28), consistent with a highly admixed population. Along the French Atlantic coast, mean Mediterranean ancestry was lower and declined gradually toward northern locations, reflecting northward diffusion of Mediterranean alleles. However, this decline was not continuous. A pronounced step was detected on the western side of the Cotentin Peninsula between SM and CH (**Fig. 2C**), where Mediterranean ancestry decreased sharply by a factor of 3.5. This pattern, previously reported in the same region (Robinet et al., 2020), marks the location of a barrier to gene flow between the BOB and NS stocks in the western Channel during summer.

### Spatio-temporal dynamics of the boundary between BOB and NS stocks

Integrating genomic data with geographic information derived from reconstructed trajectories allowed us to localize the genetic discontinuity between the BOB and NS stocks during two key phases of the sea bass life cycle. When Mediterranean ancestry was mapped using individual mean positions during the summer feeding season, the western English Channel displayed ancestry levels similar to those observed in the BOB area (**Fig. 3A**). This pattern was consistent across most of ICES subdivision 27.7.e, indicating that the stepped ancestry gradient identified from tagging locations sampled in June (**Fig. 2C**) persisted at the same position through September. In contrast, mapping Mediterranean ancestry using individual positions during the winter spawning season revealed a westward-shifted discontinuity, located slightly north of the 48th parallel, which separates the BOB and NS ICES management areas (**Fig. 3C**).

**Figure 3:**
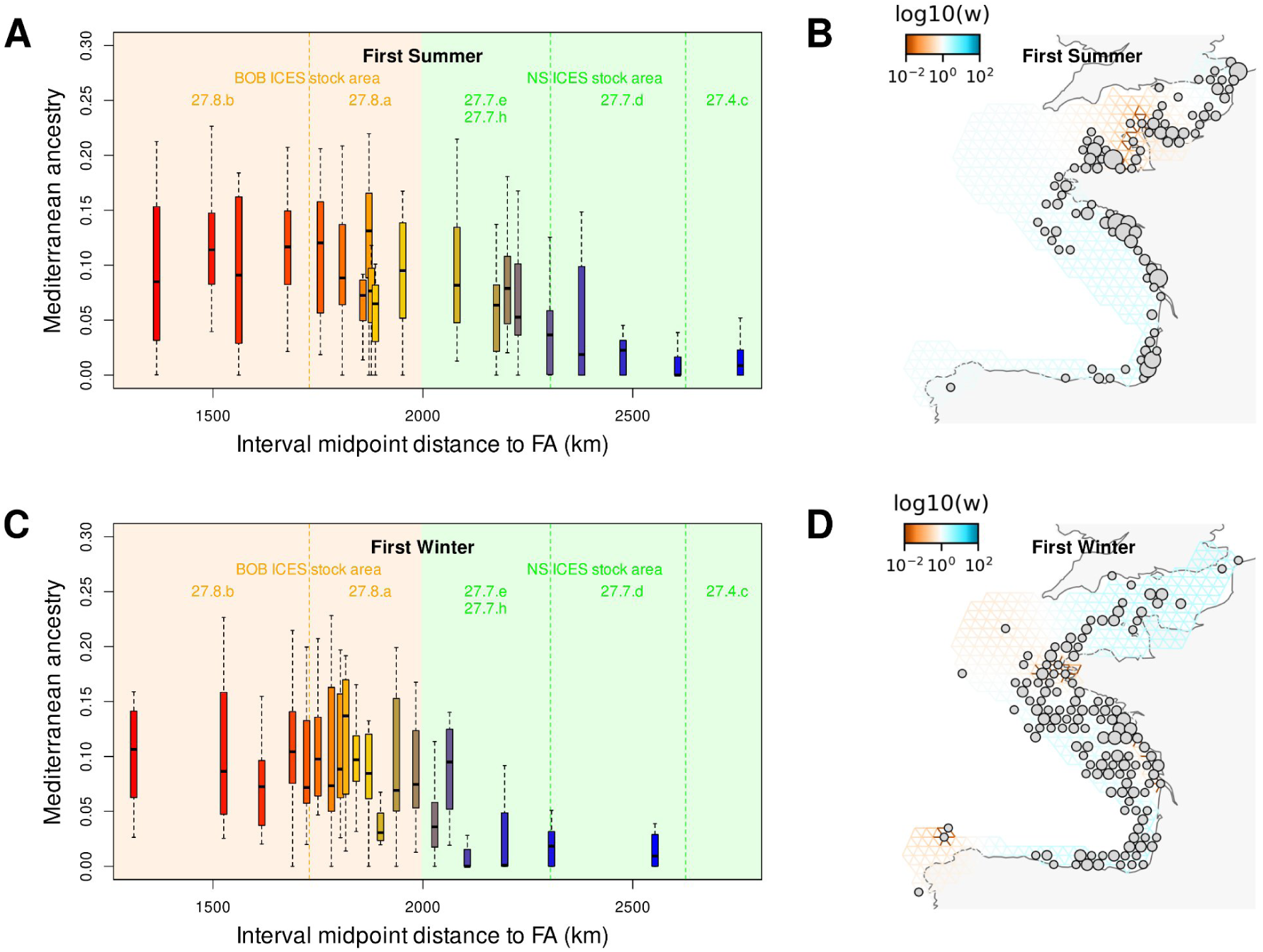
Seasonal fluctuations in the genetic discontinuity between European sea bass stocks. Integrated analysis of genetic data and reconstructed trajectories during the first summer feeding season (A, B) and the first winter spawning season (C, D) following tagging. **(A)** Summer distribution of Mediterranean ancestry, binned into 20 intervals containing equal numbers of individuals and plotted as a function of the midpoint least-cost distance to the Faro location. Mediterranean ancestry levels in the western English Channel (ICES area 27.7.e) are close to those observed in the BOB ICES area, indicating the predominance of the BOB stock during summer. **(B)** Estimated effective migration surface (EEMS) inferred from individual summer positions projected onto the nearest nodes of a dense triangular grid covering sea bass habitat. Colors represent local effective migration rates on a log_10_ scale relative to the global mean. A zone of reduced effective migration separating BOB and NS stocks is located on the western side of the Cotentin Peninsula. **(C)** Winter distribution of Mediterranean ancestry, showing a westward shift in the stepped ancestry gradient relative to summer. **(D)** Estimated effective migration surface based on individual winter spawning positions, placing the barrier of reduced effective migration between BOB and NS stocks at the tip of Brittany.

The spatio-temporal dynamics of this discontinuity was further examined using spatially explicit estimates of effective migration rates. Effective migration surfaces revealed localized barriers to gene flow at fine spatial scales, with effective migration rates up to two orders of magnitude lower than the background level. During summer, the inferred barrier was positioned on the western side of the Cotentin Peninsula, between the Channel Islands and Cap de la Hague (**Fig. 3B**). Outside this region, genetic differentiation increased only weakly with geographic distance, indicating minimal isolation-by-distance. In contrast, effective migration surfaces based on winter spawning positions identified a westward-shifted barrier near the tip of Brittany (**Fig. 3D**). Together, these analyses indicate that the genetic boundary between BOB and NS stocks is not fixed in space but shifts seasonally across the western English Channel, as feeding and spawning migrations redistribute adult sea bass between summer and winter habitats.

### Evidence for spawning-site philopatry

Tagging data have previously demonstrated spawning-site fidelity in European sea bass but could not determine whether individuals return to their natal region to reproduce (de Pontual et al., 2019; de Pontual et al., 2023; Le Luherne et al., 2022). Consequently, it has remained unclear whether regional philopatry contributes to maintaining stock structure despite temporary mixing. By integrating migration trajectories with genomic data, we found that individual spawning behavior within the English Channel area was partially associated with genetic composition. Specifically, mixed-effects modeling revealed a strong relationship between individual genetic ancestry and spawning latitude after accounting for variation among tagging locations. Individual coordinates along PC1 (from **Fig. 2B**) had a highly significant effect on winter spawning latitude (from **Fig. 1F**), with an estimated decrease of 0.427 degree in spawning latitude per unit decrease in PC1 (s.e.=0.099, p < 0.001, **Fig. 4**). Thus, within English Channel tagging locations, individuals with more BOB-like genetic ancestry tended to spawn farther south than individuals with more NS-like ancestry. This pattern is consistent with regional spawning-site philopatry. This conclusion was further supported by individuals tracked over two consecutive years, which exhibited consistent fidelity to either NS or BOB spawning areas (**Fig. 1D**). At the two tagging locations where fidelity to both spawning regions was observed (CH and DK), individuals returning to the BOB spawning area tended to be genetically more BOB-like than those returning to NS spawning areas (**Fig. 4**).

**Figure 4:**
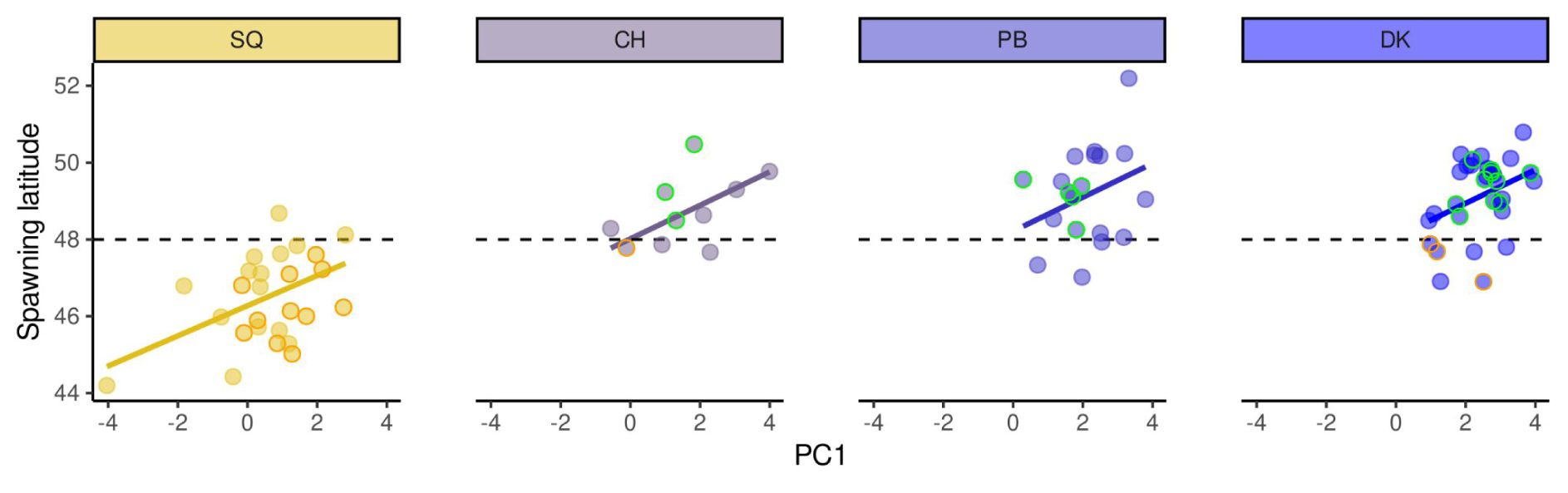
Relationship between genetic composition and spawning latitude in the NS ICES stock area. Results of a mixed-effects model relating individual genetic makeup (PC1 coordinates) to spawning-site latitude across NS tagging locations using reconstructed winter positions. Scatter plots show the relationship between individual PC1 and winter spawning latitude for each tagging location. Tagging location was included as a random intercept and slope, allowing both baseline spawning latitude and the strength of the genetic effect to vary among sites. Colored lines represent the fitted regression for each tagging location, which show similar slopes across locations. The horizontal dashed line marks the boundary between the BOB and NS ICES stock areas. Individuals exhibiting spawning-site fidelity to the BOB or NS over two consecutive winters are highlighted with orange and green circles, respectively.

Genome-wide association analysis did not identify any SNPs significantly associated with spawning latitude (**Suppl. Fig. S3**). This result suggests that spawning-site choice is unlikely to be controlled by loci with large effects, or that such effects could not be detected with the present sample size. Instead, the observed association between genetic ancestry and spawning latitude is more consistent with natal or regional homing to genetically differentiated breeding areas, rather than direct genetic control of migratory behavior.

### Recent genetic connectivity inferred from IBD sharing

Analysis of recent coancestry identified a total of 2412 identical-by-descent (IBD) segments ≥ 7 cM shared among pairs of individuals (**Fig. 5A**). The mean fraction of the genome shared IBD was significantly higher for pairs of individuals from the same tagging location than for pairs from different locations (mean kinship coefficient within locations = 2.96 × 10**⁻⁵**; between tagging locations = 2.06 × 10**⁻⁵**; *t*-test, *P* = 0.0019). Hierarchical clustering of average pairwise kinship coefficients revealed groups of recent shared ancestry that closely matched population groupings inferred from genotype-based analyses. Specifically, all tagging locations assigned to the BOB stock, including SQ, clustered together, whereas tagging locations from SM to DK formed a distinct cluster (**Fig. 5B**). These results indicate that the genetic subdivision between BOB and NS stocks is reflected in patterns of recent connectivity.

**Figure 5:**
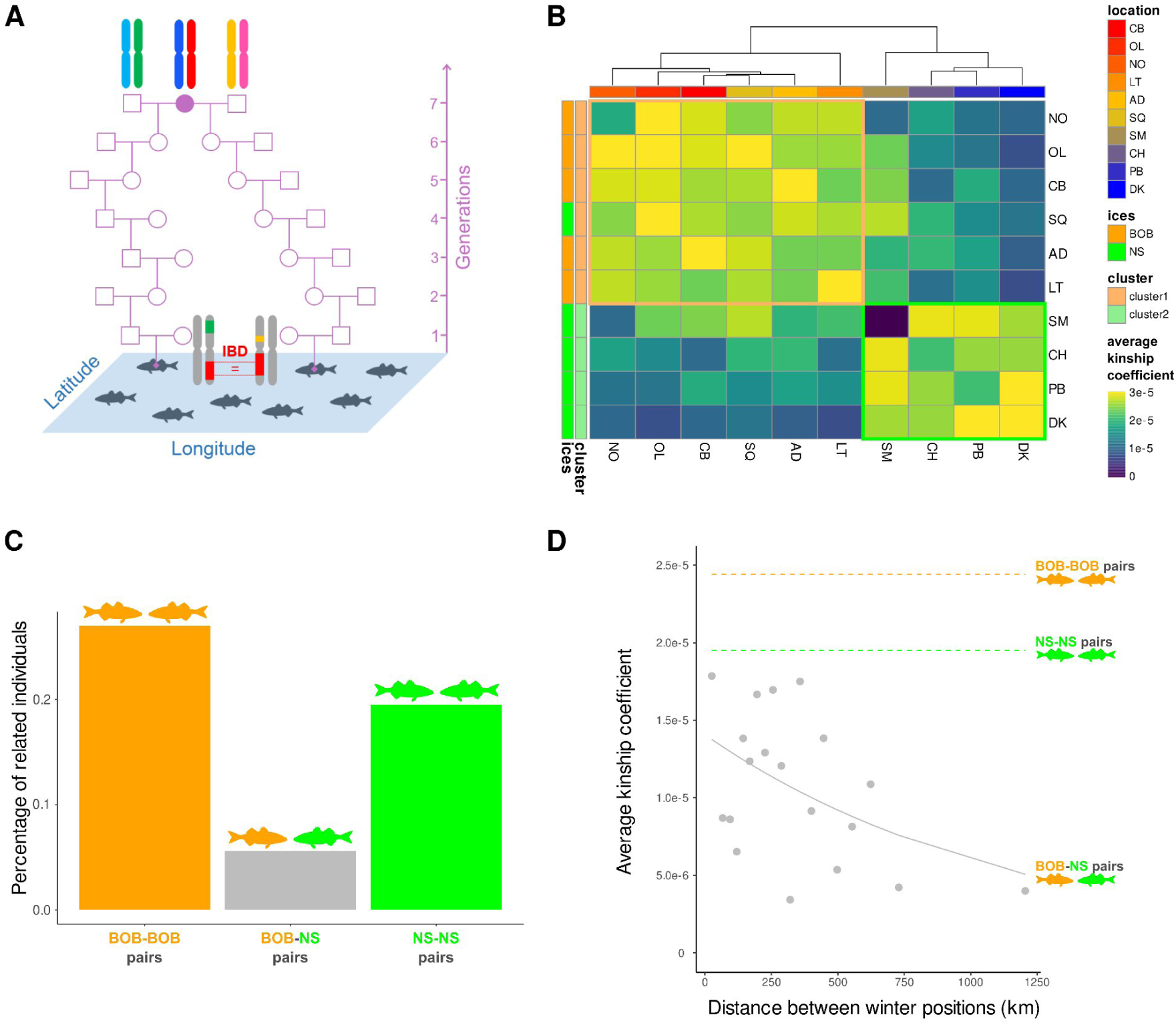
Recent genetic connectivity inferred from shared identical-by-descent (IBD) segments. **(A)** Schematic illustration of an IBD segment (red) shared between two seventh-degree relatives. IBD segments are genomic regions inherited from a recent common ancestor (female ancestor shown in purple) without intervening recombination. Because recombination progressively shortens IBD segments across generations, segment length provides information on the time since the most recent common ancestor, following a recombination-clock principle. **(B)** Heatmap of average pairwise kinship coefficients, calculated as the mean fraction of the genome shared IBD between pairs of individuals and averaged within and between tagging locations. Hierarchical clustering reveals two main clusters corresponding to the BOB and NS stocks, indicating reduced recent coancestry between stocks. **(C)** Proportion of related pairs (seventh-degree relatives or closer) within and between BOB and NS stocks, showing an approximately fourfold reduction in recent genetic exchange between stocks (BOB-NS pairs) compared to within stocks (BOB-BOB or NS-NS pairs). **(D)** Relationship between average kinship coefficient and spawning-site distance for pairs of individuals from to the same or different stocks. Points represent distance-binned mean kinship values for BOB-NS pairs, and the fitted beta regression shows a decline in relatedness with increasing spawning distance, indicating that recent inter-stock gene flow occurs primarily in the transition zone between BOB and NS spawning grounds. Horizontal dashed lines indicate mean kinship within BOB (orange) and NS (green) stocks.

Restraining the analysis to individual pairs with known spawning stock membership on reconstructed winter locations, the proportion of related pairs (seventh-degree relatives or closer) was 3.5-4.8 times lower between stocks than within stocks (**Fig. 5C**). In addition, among pairs spawning in different stocks, mean kinship coefficients declined significantly with increasing distance between spawning locations (**Fig. 5D**). Together, these results suggest that, over an approximate timescale of the last seven generations, effective migration between BOB and NS stocks is substantially lower than within stocks and primarily occurs through exchanges near the boundary between spawning areas, around the 48th parallel.

### Range-wide stock structure in the Northeast Atlantic

Integrating our dataset with published genotypes from Taylor et al. (2025) revealed coherent patterns of stock structure across the Northeast Atlantic. Sea bass sampled from water surrounding the UK predominantly aligned with the NS genetic background (**Fig. 6A**). PC1-based empirical distributions derived from individuals in our study with known spawning-stock membership showed substantial overlap between the BOB and NS reference baseline distributions, reflecting weak overall differentiation. Despite this overlap, the distribution of PC1 coordinates for individuals from Taylor’s dataset closely matched the NS reference distribution (**Fig. 6B**) with an inferred contribution of the BOB stock ranging from 0 to 0.007 (95% posterior CI).

**Figure 6.**
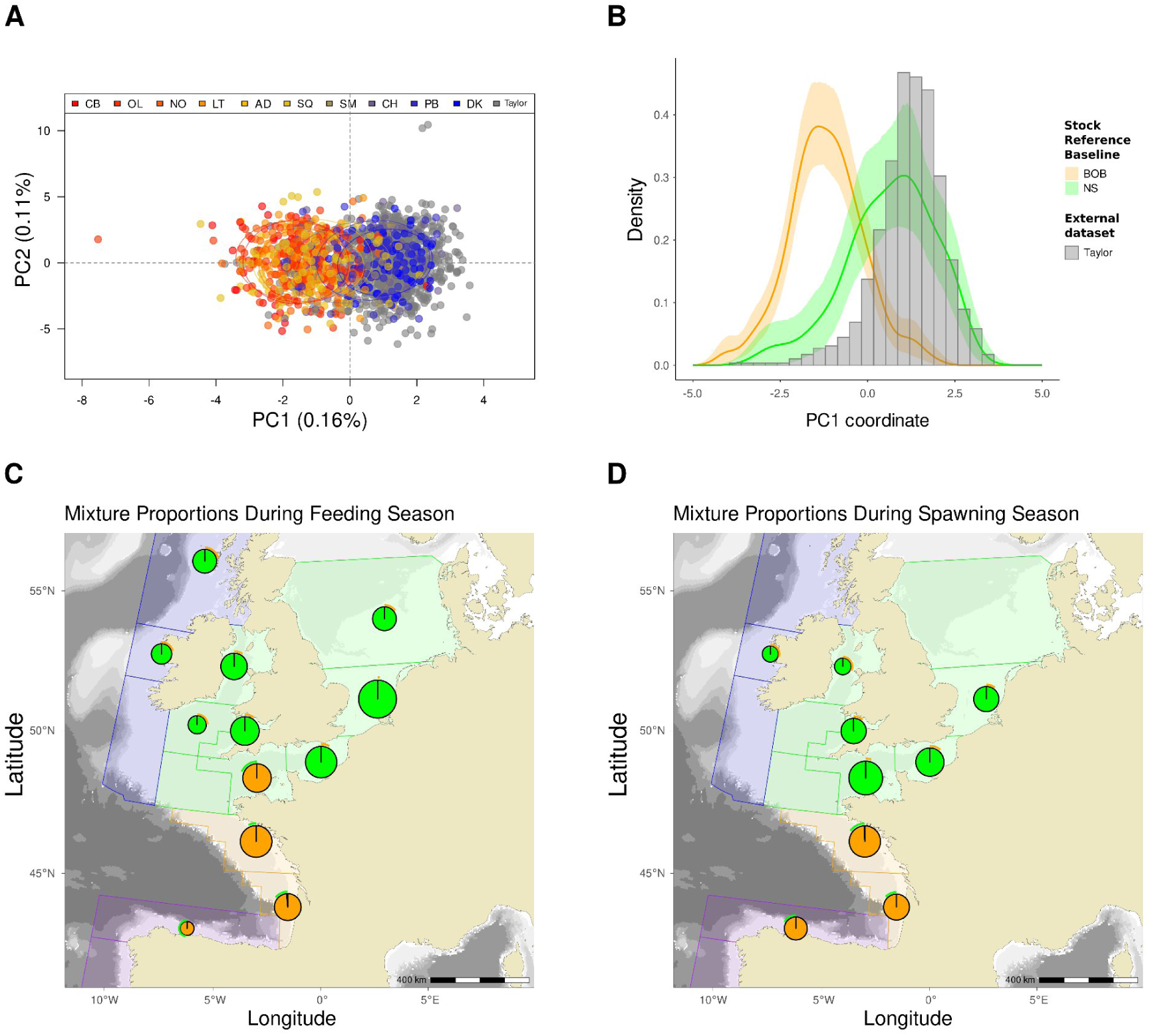
Stock structure of European sea bass in the Northeast Atlantic. **(A)** Principal Component Analysis (PCA) of the merged dataset combining individuals from this study (colored by tagging location) and from Taylor et al. (grey), showing the distribution of sample coordinates across the Northeast Atlantic, along the PC1 axis reflecting the gradient of Mediterranean ancestry. **(B)** Empirical densities of PC1 coordinates for individuals assigned to the BOB (orange) and NS (green) stocks using tagging data (baseline reference distributions), compared with the histogram distribution (grey) of individuals from the Taylor et al. dataset. Overlap between BOB and NS baseline distributions reflects weak genetic differentiation, while samples from Taylor et al. (2025) align predominantly with the NS distribution. **(C)** Map of the spatial distribution of inferred BOB stock proportions across ICES areas during the feeding season. Pie charts represent the estimated proportion of BOB (orange) versus NS (green) individuals in each area, highlighting a major contribution of BOB in southern areas (27.8) and the western English Channel (27.7.e), and a major contribution of NS in more northern regions. **(D)** Spatial distribution of inferred BOB stock proportions across ICES areas during the spawning season. The major contribution of BOB is maintained in southern areas (27.8), whereas areas within division 27.7 (including 27.7.e) show near-zero BOB contribution, indicating NS dominance north to the 48th parallel. Confidence intervals are shown as outer colored arcs representing the 95% CI of the minor stock in each area. Larger uncertainty is associated with areas of smaller sample size, as shown by the radius of circles representing the squarerooted sample size.

Mixture analyses revealed marked spatial structuring during both feeding and spawning periods. During the feeding season, areas 27.8.c, 27.8.b, 27.8.a, and 27.7.e were almost exclusively dominated by the BOB stock (inferred BOB proportion ≈ 1; **Fig. 6C**). In contrast, other areas within division 27.7 (g, f, d, b, a) and more northern regions (27.6.a, 27.4.b–c, 27.3.a.20) were almost entirely composed of the NS stock. During the spawning season, spatial segregation was shifted southward: areas 27.8.c, 27.8.b, and 27.8.a remained dominated by the BOB stock, whereas all areas within division 27.7 (including 27.7.e) showed near-zero BOB contribution, indicating NS dominance up to the 48th parallel (**Fig. 6D**).

Confidence intervals were generally narrow across areas in both seasons (mean width of 95% posterior CI: 0.17 in feeding season, 0.16 in spawning season), with wider intervals mainly associated with area of small sample sizes. Resampling of reference individuals yielded highly consistent assignments, indicating low sensitivity of the estimates to baseline composition (**Suppl. Table S2**). Overall, these results support the presence of two weakly differentiated but demographically structured sea bass stocks occupying distinct ICES divisions, with limited mixing at their boundary.

## Discussion

Understanding the stock structure of European sea bass has long challenged both movement ecology and population genetics, limiting the translation of scientific knowledge into clear management guidance. These difficulties stem from extensive seasonal movements, weak genetic differentiation, and diverse migratory strategies across the species’ range. By integrating high-density SNP genotyping with individual movement data across seasons, we characterize the spatio-temporal dynamics of a shifting connectivity boundary between the Northern (NS) and Bay of Biscay (BOB) stocks. In winter, this boundary aligns with the current stock delineation near the 48th parallel, whereas in summer it shifts northeastward to the west of the Cotentin Peninsula. Despite seasonal mixing in the English Channel, regional spawning-site philopatry to genetically differentiated breeding areas appears sufficient to maintain demographic independence between stocks. This conclusion is supported by analyses of recent genetic ancestry based on IBD segments, which show reduced sharing of recent common ancestors between stocks relative to within stocks. Together, these results indicate that the two evidenced subpopulations experience limited contemporary connectivity, with direct implications for the monitoring and management of European sea bass in the Northeast Atlantic.

### Seasonal migrations of European sea bass

Behavioural profiles and migration strategies from tagging studies have revealed a partially migratory meta-population structure in European sea bass. While juveniles are largely sedentary (Stamp et al., 2021), adults either remain resident in coastal areas (de Pontual et al., 2019; Doyle et al., 2017; O’Neill et al., 2018) or undertake seasonal migrations between inshore feeding habitats and offshore spawning grounds, often exhibiting strong fidelity to both summer feeding and winter spawning sites (de Pontual et al., 2019; de Pontual et al., 2023; Goossens et al., 2024; Le Luherne et al., 2022; Wright et al., 2024). Consistent with these observations, our results indicate that long-distance migrations are more frequent in the NS area (**Fig. 1B**), suggesting stronger environmental constraints toward the northern edge of the species’ distribution.

Tagging studies from the Celtic Sea, English Channel, and North Sea identified the western English Channel as a dynamic interface between the NS and BOB ICES stocks, characterized by diverse migration strategies. During the spawning season, individuals from the English Channel and western UK coasts are joined by migrants from the southern North Sea (Pickett et al., 2004; Wright et al., 2024). Sea bass tagged in the southern North Sea typically migrate southwest in winter toward the English Channel and the Celtic Sea, although some remain locally, suggesting possible spawning within the southern North Sea (Goossens et al., 2024).

Tagging along the French coast further revealed marked regional differences within the English Channel. Individuals tagged in the eastern Channel predominantly spawn within the Channel or the Celtic Sea, and only occasionally in the northern Bay of Biscay (Bertignac et al., 2026; de Pontual et al., 2019; de Pontual et al., 2023). In contrast, individuals tagged in summer in the western Channel or Iroise Sea primarily migrate south to spawn in the Bay of Biscay (de Pontual et al., 2023, **Fig. 1E,F**). Together, these patterns indicate transient mixing between NS and BOB stocks in the western English Channel, while spawning- and feeding-site fidelity contribute to partial spatial segregation across seasons.

### Evidence for a seasonally shifting stock boundary

Using an independent set of samples and genetic markers, our study identifies the same genetic discontinuity between NS and BOB stocks as previously reported by Robinet et al. (2020). This pattern is first evident in the hierarchical clustering of tagging locations (**Fig. 2A**), which groups eastern English Channel locations into a distinct cluster showing weak but significant genetic differentiation (*F*_ST_ ≈ 0.0005) from BOB locations. This signal becomes more pronounced when genetic structure is examined through variation in Mediterranean ancestry (**Fig. 2B,C**), revealing the same sharp transition previously described in the Saint-Malo region (Robinet et al., 2020).

Previous works have shown that alleles of Mediterranean origin tend to be deleterious when introgressed into Atlantic genomes (Duranton et al., 2018, 2020; Tine et al., 2014). Although Mediterranean ancestry continuously diffuses northward through introgression, it is progressively purged by selection, generating a latitudinal gradient along the Northeast Atlantic coast (Robinet et al., 2020). Local reductions in gene flow can therefore produce sharp transitions in this gradient, maintained by a balance between introgression and selection.

Our analysis of Mediterranean ancestry as a function of distance from southern Portugal shows that the position of the ancestry step shifts seasonally, from the southern boundary of ICES division 27.7.e in winter to its western margin in summer (**Fig. 3A,C**). This temporal displacement likely reflects large-scale seasonal movements of sea bass throughout their annual cycle, as reported in other marine teleosts and elasmobranchs in the same area (Griffiths et al., 2020; Horton et al., 2025; Hunter et al., 2004). Rather than eroding the winter discontinuity through mixing, feeding migrations thus appear to translate it northeastward across the western English Channel (**Fig. 3B,D**).

### Regional philopatry maintains demographic independence between stocks

Within the narrow transition zone between BOB and NS stocks, effective migration rates are two orders of magnitude lower than within stocks, indicating markedly reduced genetic connectivity. Interpreting this pattern requires considering the mechanisms underlying spawning-site fidelity. Although these mechanisms remain poorly understood in European sea bass, several non-exclusive processes may be involved. Fidelity may arise from learned environmental preferences or social processes, such as cultural transmission (Meager et al., 2018), as well as from genetically influenced behaviours such as natal homing. Otolith microchemistry analyses have not provided evidence for natal homing in sea bass (Le Luherne et al., 2022). If fidelity were driven by strict natal homing, one would expect temporally mixing but demographically independent stocks. However, the partial spawning-site fidelity observed in our data (**Fig. 1C,D**) indicates that exchanges between stocks do occur. Even limited exchanges concentrated in the transition zone between stocks may generate sufficient gene flow to explain the weak genetic differentiation reported in previous studies (Coscia & Mariani, 2011; Fritsch et al., 2007; Robinet et al., 2020; Souche et al., 2015). This highlights the need to combine movement and genetic data to determine whether early-life dispersal and partial adult homing are sufficient to ensure demographic coupling between stocks.

To address this, we directly link long-term genetic structure with recent demographic connectivity. Our results provide direct evidence of regional spawning-site philopatry (**Fig. 4**), indicating that reproductive homing is the primary mechanism maintaining genetic differentiation between stocks. Using IBD segments, we characterize cryptic relatedness up to approximately seven generations back in the population pedigree (**Fig. 5A**). Patterns of recent relatedness recapitulate the stock structure observed at longer timescales (**Fig. 5B**) and show that contemporary connectivity between stocks is approximately four-fold lower than within stocks (**Fig. 5C**). Moreover, recent kinship between stocks is concentrated around the transition zone, consistent with localized exchange across the stock boundary. Overall, these findings indicate that exchanges due to partial spawning-site fidelity occur at rates too low to ensure demographic connectivity between the BOB and NS stocks. Stock mixing thus likely represents a transient phase between periods of spatial segregation in the sea bass annual cycle.

### Limits and future of the combined approach

Our study has limitations inherent to both tagging and genetic approaches. The accuracy and precision of reconstructed fish tracks depend on the geolocation model and the quality of environmental data. Simulation-based sensitivity analyses have demonstrated the reliability of the sea bass geolocation model in the Iroise Sea (Woillez et al., 2016), but its performance may vary across the broader range of migration behaviours encountered in our large-scale survey.

To address this, model adaptations were implemented to capture behavioural diversity, including attraction to thermal plumes, switching between low- and high-activity movement states, and the use of multi-layer temperature likelihoods (de Pontual et al., 2023). Although several validation approaches have been proposed (Gatti et al., 2021; Haase et al., 2021), they remain poorly suited to the spatial and temporal scales considered here. Current geolocation models typically achieve average errors of ∼30-50 km for demersal fish and ∼120 km for large pelagic species (Gatti et al., 2021), which are likely sufficient to detect broad barriers to gene flow. However, winter reconstructions may be less reliable than summer estimates due to greater temporal distance from tagging and weaker thermal gradients. Integrating ground-truth data from double-tagging approaches, combining acoustic and DST archival tags, offers a promising avenue to improve model validation and track reconstruction (Gonse et al., 2025; Goossens et al., 2023; Liu et al., 2017). Despite these limitations, the consistency of our results suggests that combining genetic and tagging data provides sufficient resolution to detect fine-scale spatio-temporal connectivity patterns in European sea bass.

On the genetic side, inference of recent coancestry may be affected by false positives or missed detections of IBD segments, although these risks were minimized through standard filtering procedures. Beyond these limitations, IBD segments are likely to become a powerful source of information for genetic stock identification based on ancestry sharing. Because the number of shared ancestors increases rapidly with genealogical depth, recent coancestry can be detected even in the absence of close-kin pairs. This approach extends inference beyond the first and second degrees of relatedness typically evaluated in close-kin mark-recapture (CKMR) studies (Bravington et al., 2016; Trenkel et al., 2022), without requiring very large sample sizes.

In parallel, traditional individual assignment methods based on genetic similarity (Chen et al., 2018; Kuismin et al., 2020; Moran & Anderson, 2019) have limited power in sea bass due to weak genetic differentiation. Rather than assigning individuals, we estimated stock composition using a likelihood-based mixture model, which avoids misclassification in the presence of overlapping reference distributions and provides estimates of stock proportions with explicit measures of uncertainty. These sample-level mixture proportions directly correspond to the parameters of interest for stock assessment. However, our inference relied on summarizing individual genetic variation through an ancestry-oriented PCA obtained with genotypes, and could benefit from emerging approaches such as branch PCA (Lehmann et al., 2026), which better capture recent population structure over specific time intervals by leveraging information from recent IBD segments.

### Implications for management

Individual movement shapes seasonal population dynamics and is therefore a key component of management in partially migratory species (Reid et al., 2018). In European sea bass, seasonal migrations interact with life-history traits such as longevity (>20 years), late maturity (4-6 years), spawning aggregation and fidelity to localised foraging areas (Doyle et al., 2017), increasing vulnerability to overexploitation. Fishing pressure on demographically independent stocks can therefore lead to rapid local depletion, even when overall abundance remains above precautionary thresholds (Erisman et al., 2011). This mechanism may have contributed to the recent decline of the NS stock, which has shown partial recovery following management measures implemented since 2015 (ICES, 2025b).

Sustainable fisheries management relies on accurate identification of biologically meaningful stock units (Kerr et al., 2017; Seljestad et al., 2024). Our range-wide analysis provides clear evidence that European sea bass in the Northeast Atlantic comprise two genetically distinct stocks, the Bay of Biscay (BOB) and Northern (NS) stocks, in addition to the western Iberian stock not included here. Regions from West Ireland and Scotland were not sampled but are likely part of the NS stock, extending its northern range. By integrating genomic and tagging data, we reconcile previous discrepancies among studies limited in seasonal resolution (Robinet et al., 2020), spatial coverage (Taylor et al., 2025), or genomic representation (Fritsch et al., 2007; Souche et al., 2015). In particular, the genetic discontinuity observed between the eastern and western English Channel (Robinet et al., 2020) reflects the summer configuration of stock structure (**Fig. 6C**), whereas its absence in Taylor et al. (2025) results from sampling restricted to the NS stock (**Fig. 6A,B**). Samples from ICES area 27.7.e in Taylor et al. (2025) were collected only during the spawning season, and the absence of sampling during the feeding season in the western English Channel precluded detection of the summer expansion of the BOB stock and its potential mixing with the NS stock. This highlights the importance of comprehensive seasonal sampling when identifying stock units and interpreting population structure.

A key question is whether genetic connectivity reflected by weak genetic differentiation masks demographic independence between the BOB and NS stocks. Our results show that partial spawning-site fidelity leads to localized exchanges near the transition zone at the tip of Brittany, but these remain limited in extent and occur at much lower rates than within-stock exchanges (**Fig. 5C,D**). This restricted connectivity indicates that exploitation of one stock cannot be compensated by immigration from the other, increasing the risk of local depletion and reinforcing the need for stock-specific management.

The spatio-temporal dynamics of stock structure emerge as a key element for European sea bass fisheries management. Spatial genetic analyses and mixture proportion maps show that the boundary between BOB and NS stocks shifts between winter and summer, reflecting large-scale seasonal migrations. Tagging data broadly support these dynamics but also highlight uncertainties related to sampling biases. For example, the apparent absence of southward spawning migrations in UK fish tagged in the western Channel likely reflects late-autumn tagging of NS individuals (Wright et al., 2024), whereas summer tagging along the French coast and Channel Islands indicates movement toward the Bay of Biscay (de Pontual et al., 2023; Pawson et al., 2007). These findings, consistent with our estimates of seasonal stock composition in the western English Channel (**Fig. 6C,D**), emphasize the need to interpret tagging data within a seasonal framework.

Delineation of stock boundaries often follows jurisdictional lines that do not align with biological units, potentially leading to mismatches in management (Cadrin et al., 2023). Our estimates of mixture proportions indicate that the current ICES boundary at the 48th parallel provides an effective delineation of BOB and NS stocks during winter. In area 27.7.e, the genetic composition strongly supports the predominance of NS individuals during the spawning season, with an upper confidence bound indicating at most 4.6% BOB contribution (**Fig. 6D**).

In contrast, the summer configuration, where the BOB stock appears to dominate, is less precisely resolved due to smaller sample sizes and wider confidence intervals (up to 18.4% NS contribution). This uncertainty is compounded by the spatial restriction of sampling to the French side of the western English Channel, highlighting the need for more balanced spatial sampling during the feeding season. If confirmed by finer-scale analyses, the boundary between areas 27.7.e and 27.7.d could better reflect biological structure during summer.

Taken together, these results call for a refinement of current management practices to account for the dynamic nature of stock structure. First, stock assessment and catch reporting in area 27.7.e should explicitly incorporate seasonal variation in stock composition, ideally distinguishing between French and UK waters where relative stock proportions may differ. Second, enhanced seasonal sampling in the western English Channel is needed to better quantify stock mixing and reduce uncertainty in mixture estimates by improving the genetic baseline. Finally, integrating genetic mixture models with tagging data into stock assessment frameworks would allow explicit estimation of seasonal exchange rates between stocks (Bertignac et al., 2026), improving the biological realism of management models.

## Conclusion

Our study highlights the value of combining tagging and genetic data to understand how migratory connectivity and partial migration shape the spatio-temporal dynamics of population structure and demography (Müller et al., 2023; Reid et al., 2018; Webster et al., 2002). Recent advances in genome-wide genealogy inference (Lewanski et al., 2024; Nielsen et al., 2025) will soon make it possible to link individual genetic ancestry with recent population pedigree and movement data. This integration will bridge temporal gaps between long-term and contemporary measures of connectivity, while strengthening links between genetic and demographic concepts. By leveraging genealogical frameworks, these approaches offer new opportunities to derive quantitative estimates of key demographic parameters relevant to conservation and management.

## Data Accessibility

Original data used in this study are publicly available on SEANOE: https://doi.org/10.17882/117393. Scripts to reproduce the figures are available on GitHub: https://github.com/PA-GAGNAIRE/Integrating-genomic-and-tagging-data-in-sea-bass.git

## Acknowledgements

This study was part of the *Barfray* project funded by the European Maritime and Fisheries Fund (EMFF-OSIRIS N°: PFEA 400017DM0720006), France Filière Pêche (FFP), the French Ministry of Agriculture and Food (MAF) and IFREMER. The DST data are from the national *BarGip* project (funded by IFREMER, MAF and FFP). We thank Rubén Lechuga for providing samples from the Algarve region in southern Portugal. We thank the Gentyane platform (INRAe, Clermont-Ferrand) for the genotyping.

