## Supplementary Materials file for "Integrating genomic and tagging data reveals spatio-temporal population structure in Northeast Atlantic European sea bass"

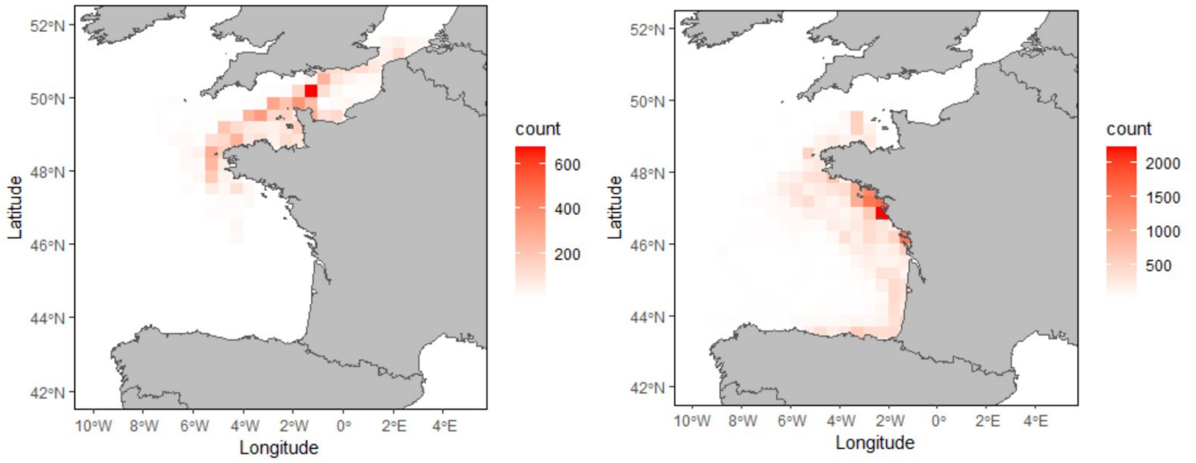


**Figure S1:** Spatial distribution of tagged sea bass grouped according to the main spawning migration strategy of the tagging sites over the first winter semester after tagging. Left: Count per block of trajectory points (i.e. days) from individuals belonging to DK, PB and SV with NS spawning migration strategy. Right: Count per block of trajectory points (i.e. days) from individuals belonging to SM, SQ, AD, LT, NO, IO and CB with the BOB spawning migration strategy.

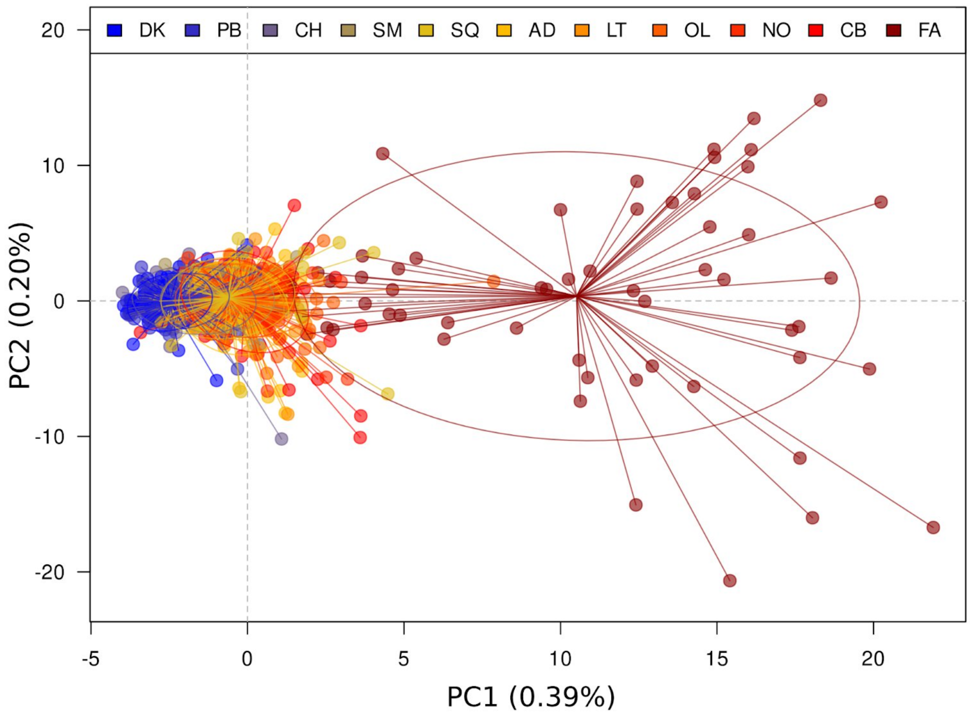


**Figure S2:** Principal Component Analysis of northeast Atlantic populations of European sea bass, including all individuals (N=765) from the 10 French Atlantic tagging locations and southern Portugal.

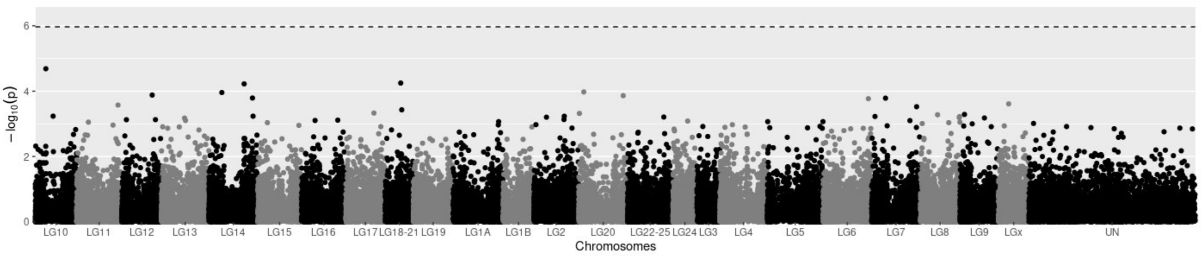


**Figure S3:** GWAS testing SNP associations with individual spawning latitude, based on N=248 individual mean latitudinal positions during the first winter. The dashed horizontal line represents the genome-wide significance threshold, adjusted using Bonferroni correction with an alpha of 0.05.

**Supplementary Table S1. Summary of the 10 tagging surveys conducted along the French Atlantic and English Channel coasts.** Survey codes are assigned to sampling locations, listed here from north to south.

| Survey Code (Date) | Location | Number of Tagged fish | Number of Recoveries | Number of constructed tracks |
|---:|---:|---:|---:|---:|
| DK (3-13 June 2014) | Dunkerque | 150 | 57 | 48 |
| PB (1-12 June 2015) | Port-en-Bessin | 89 | 39 | 29 |
| CH (21-24 June 2016) | Saint-Vaast-la-Hougue | 129 | 37 | 25 |
| SM (9-11 Sept. 2015) | Saint-Malo | 16 | 5 | 1 |
| SQ (17-27 June 2014) | Saint-Quay-Portrieux | 152 | 65 | 41 |
| AD (9-19 June 2015) | Audierne | 179 | 66 | 49 |
| LT (9-19 Sept. 2014) | La Turballe | 143 | 70 | 50 |
| NO (31 Aug. - 4 Sept. 2016) | Noirmoutier | 107 | 43 | 35 |
| OL (7-16 Sept. 2015) | Ile d’Oléron | 132 | 41 | 36 |
| CB (1-11 Sept. 2014) | Capbreton | 123 | 59 | 50 |
| | **Total DSTs** | **1220** | **482** | **364** |

**Supplementary Table S2. Inferred mixture proportions within ICES areas during (A) FEEDING and (B) SPAWNING seasons.** Within each ICES area, Feeding and Spawning samples from Taylor et al (2025) were joined with the samples from this study based on their Summer and Winter reconstructed position. N: sample size in each area, BOB_Pi: Maximum *a posteriori* estimate proportion of the BOB stock in the mixture sample, along with it posterior 95% credible intervals (BOB_Pi_low, BOB_Pi_hi). BOB_Pi_boot: Maximum *a posteriori* estimate proportion of the BOB stock in the mixture sample after resampling reference individuals with replacement across 1000 bootstrap replicates, along with 95% percentile intervals derived from the distribution of bootstrapped estimates (BOB_Pi_boot_low, BOB_Pi_boot_hi).

### A. Inferred mixture proportions during FEEDING season:

| Area_Full | N | BOB_Pi | BOB_Pi_low | BOB_Pi_hi | BOB_Pi_boot | BOB_Pi_boot_low | BOB_Pi_boot_hi |
|---|---|---|---|---|---|---|---|
| 27.8.c | 7 | 1.0000 | 0.5265 | 0.9960 | 1.0000 | 1.0000 | 1.0000 |
| 27.8.b | 54 | 0.9870 | 0.8579 | 0.9990 | 0.9848 | 0.9670 | 1.0000 |
| 27.8.a | 113 | 1.0000 | 0.9309 | 1.0000 | 0.9999 | 1.0000 | 1.0000 |
| 27.7.e | 70 | 1.0000 | 0.8138 | 0.9980 | 0.9827 | 0.9179 | 1.0000 |
| 27.7.g | 14 | 0.0000 | 0.0010 | 0.2422 | 0.0000 | 0.0000 | 0.0000 |
| 27.7.f | 71 | 0.0000 | 0.0000 | 0.0881 | 0.0005 | 0.0000 | 0.0030 |
| 27.7.d | 109 | 0.0000 | 0.0000 | 0.0751 | 0.0056 | 0.0000 | 0.0430 |
| 27.7.b | 20 | 0.0000 | 0.0010 | 0.2062 | 0.0009 | 0.0000 | 0.0100 |
| 27.7.a | 51 | 0.0000 | 0.0000 | 0.0941 | 0.0000 | 0.0000 | 0.0000 |
| 27.6.a | 34 | 0.0000 | 0.0020 | 0.2162 | 0.0079 | 0.0000 | 0.0771 |
| 27.4.c | 285 | 0.0000 | 0.0000 | 0.0170 | 0.0000 | 0.0000 | 0.0000 |
| 27.4.b | 34 | 0.0000 | 0.0010 | 0.1471 | 0.0026 | 0.0000 | 0.0360 |
| 27.3.a.20 | 14 | 0.0000 | 0.0020 | 0.2863 | 0.0001 | 0.0000 | 0.0000 |

### B. Inferred mixture proportions during SPAWNING season:

| Area_Full | N | BOB_Pi | BOB_Pi_low | BOB_Pi_hi | BOB_Pi_boot | BOB_Pi_boot_low | BOB_Pi_boot_hi |
|---|---|---|---|---|---|---|---|
| 27.8.c | 30 | 1.0000 | 0.8328 | 0.9990 | 1.0000 | 1.0000 | 1.0000 |
| 27.8.b | 47 | 1.0000 | 0.8719 | 1.0000 | 1.0000 | 1.0000 | 1.0000 |
| 27.8.a | 106 | 0.9930 | 0.8509 | 0.9980 | 0.9739 | 0.9149 | 1.0000 |
| 27.7.e | 154 | 0.0000 | 0.0000 | 0.0460 | 0.0028 | 0.0000 | 0.0260 |
| 27.7.f | 43 | 0.0000 | 0.0000 | 0.1091 | 0.0000 | 0.0000 | 0.0000 |
| 27.7.d | 67 | 0.0000 | 0.0000 | 0.1131 | 0.0082 | 0.0000 | 0.0591 |
| 27.7.b | 10 | 0.0000 | 0.0020 | 0.2993 | 0.0000 | 0.0000 | 0.0000 |
| 27.7.a | 11 | 0.0000 | 0.0020 | 0.3253 | 0.0000 | 0.0000 | 0.0000 |
| 27.4.c | 42 | 0.0000 | 0.0000 | 0.1011 | 0.0000 | 0.0000 | 0.0000 |
